# Machine Learning connects structure, bitterness & mechanism to antimalarial activity

**DOI:** 10.1101/2023.09.08.556833

**Authors:** S Sambu

## Abstract

Machine learning allows us to identify patterns we might otherwise have missed in the data. It therefore provides an ample solution to the age-old problem of lossless research. When the pattern is robust and the signal is durable, it is possible to use relatively modest amounts of data to build powerful well-learned models. By constructing a classifier using gradient-boosted machines, it was shown that the model was robust, yielded high quality metrics and demonstrated consistent probability assignments for chemistries that have a suitable chemical backbone qualified for inclusion in antiplasmodic libraries. Critical to model development was the utilization of molecular fingerprinting and extraction of physico-chemical parameters relevant to the mechanism driving antiplasmodic activity. Subsequently, such an approach allows the model to uncover the link between bitterness, molecular structure and therapeutic value. This approach to evaluating antiplasmodic activity in chemistries provides a low-cost tool usable in identifying new classes of molecules for use in reducing malaria morbidity that often affects vulnerable members of the community the most. Additionally, given their relatively broad, low colligation and potent efficacies, these molecules may provide strong safety margins and durability netting high returns for health equity.

## 1 Introduction

Development of new medicines is often compared to an arms race between the evolution of the pathogen towards drug-resistant forms and the development of new drugs. However, it can be a long, expensive and often failure-prone process(1) despite the advent of automated, multiplexed wet lab techniques and rule-driven preselection of development candidates/targets(2). One potential approach to reducing both time, costs and error-rates during drug development is the application of machine learning(3) to uncover novel connections inspired by classical chemometrics.

However, while classical chemometrics are human-readable and interpretable, being products of natural intelligence, machine learning (ML) is often not human readable, and its interpretation does not lend itself to crystallization into neat axiomatic rules for further querying, development and application. The interpretability of ML is by admission an after-thought given the earliest objectives for ML were the automation of tasks that were deemed to be narrow in scope and of limited aetiological interest for example the classification of images for machine vision. These contests were nevertheless interesting computing challenges. It is no wonder therefore, that many ML algorithms are black-box models with little mechanistic depth attributable to their success.

If ML is to be useful in a drug discovery program, there is a need to go beyond the initial mathematical horizon of minimizing error-rates. There is a need to challenge the objective setting to include the selection of a mechanistically optimal solution over a competing, yet abstract, black-box model. To this end, there have been attempts to develop post-hoc annotation routines in the form of heat maps over molecular structures(4). However, even these tend to be better used as validation modules than discovery-and-enrichment modules. There are few axiomatic principles applicable(5) especially given the diverse structures necessary for the development of a powerful algorithm. Therefore, the in-training design needs to change to allow the algorithm to display an evolution towards mechanistically-sound principles with mathematical precepts.

Molecular fingerprints have been developed as pithy inputs to the development of extensible structure-based models for predicting activity(6). In some cases, a direct prediction is favoured for computational efficiency and amenability to the often data-hungry machine learning routines such as deep learning(7). These methods require further development in their mechanistic verification given they are black-box models(6). A potential approach may be to build into the algorithm selection procedures that can find complementarity between structure and mechanism during the training phase of the model development. However, it must not stop there, rather, the algorithm needs to find a fitness benefit attributable to the mechanism discoverable in the input space that the structural input space could not provide.

Inbuilt mechanistic favour within the ML model may enable the model to better bridge across training set boundaries since structure and/or mechanism may overlap with similar models. Where the stated objective in drug development is the durability of the solution against an evolving pathogenic landscape, this agility may help to unite both preventative/prophylactic (primary health) goals and curative (secondary health) goals. Unearthing molecules that can be found within the nutritional landscape but have appreciable prophylactic (if therapeutic) purposes may help develop a multifaceted platform solution to drug development.

Although nutrition is appreciated as a cornerstone of public health, it is largely ignored in drug development despite their patient-linked synergistic relationship(8). It can be argued that gustation, a product of the long-evolved human sensory system, is the link between the two. Specifically, bitterness encodes a molecular hash whose contents relate a molecular key to physiologic values. The two most important physiologic values are toxicity and therapeutic value(9–11). Hence, were we to unlock the algorithmic mechanistic causeway between molecular structure and physiologic values, there may be a new screening platform for preselection routines and rulesets for molecular redesign towards greater potency with a margin of safety for potential drug resistance (durability).

Molecular candidate affinity at the intersection of nutrition, prophylaxis and therapy has previously been identified amongst some human communities(12). The desire and tolerance for bitter molecules amongst humans living in areas that are endemic has been linked to the presence of sub-toxic and therapeutic levels of cyanogenic species which consequently result in lower levels of parasitaemia(12). These findings underscore the importance of linking molecular structure to their corresponding taste, safety and therapeutic values during a drug design workflow.

These referenced studies have previously demonstrated the power of ML to link structure to activity directly and the limitations of interpreting said models. Moreover, the development of psychophysical models of bitterness and their linkage to compliance, safety and therapeutic potential have also been demonstrated. This article looks to provide an approach that introduces mechanistic insight into the selection process while retaining the structural basis for both the mechanism and the model’s convergence. Ultimately, as a post-hoc model of conceptual verification, the probabilistic assignments of the model are matched to independent bitterness quotients to investigate the structure-mechanism link between psychophysical neighbourhoods and the predicted antimalarial activity.

## 2 Materials and Methods

### 2.1 Metadata Collection

Data was collected from inhibition assay data deposited on PubChem as a foundational compound screening assay for antiplasmodic activity. The assay uses lactate dehydrogenase as a marker for growth of plasmodia. Plasmodia were exposed to the test compounds, with DMSO as a positive control. A 1:1 chloroquine-artemisinin mixture was used as a negative control. The dataset contains 13,533 compounds with 58.5% being active and 40.6% being inactive and the rest (0.9%) being unspecified.

### 2.2 Computational techniques for the generation of molecular descriptors

#### Variable Generation

Molecular SMILES were generated from structure identifiers (SID) using a PubChem Database and then checked for consistency.

#### Variable Transformation

Molecular SMILES were imported into R and transformed into their text-based chemical table files containing atom coordinates and corresponding bonds. These were used to generate descriptors using R Chemoinformatic libraries: Rcpi(13), ChemmineR(14) and ChemmineOB(15). Descriptors include compact standard molecular fingerprint, mass, Hydrophobicity and electrotopological descriptors. To confirm usability of the data, a random forest model was used to evaluate variable importance. See Table 2 for the descriptor set.

#### Chemical Scaffold Determination

The chemical scaffolds were extracted using clustering techniques implemented in Scaffold Hunter(16). Briefly, chemical identifiers, SMILES and antiplasmodic activity were imported into an HSQLDB database. From the database, scaffold clouds and tree maps were constructed to represent the structural hierarchy and contributions respectively within the antiplasmodic molecular library. Chemical scaffolds are clustered via a sequential agglomerative hierarchical non-overlapping clustering (SAHN) algorithm(17,18). Scaffolds were obtained by using a reductive routine that prunes lower-level branches to unearth a cipher of communal fundamental molecular substructures.

#### Model Development

The model development had 4 distinct stages. Initially, the molecular smiles were interpreted into corresponding structures represented in bit fingerprints (FP). The second involved assigning physico-chemical (PC) descriptors to the structures. A third step generated a dimensionally reduced, high-variance matrix of the molecules which captured the input as a fingerprint-physicochemical (FP-PC) space. This step removed the low magnitude vectors and zero-variance vectors in the FP-PC space. The fourth step involved developing a gradient boosting model from the FP-PC scope (GBM-FP-PC) of increasing fitness with each cycle to regress the antiplasmodic activity class assignment against the input space for the training set. The stopping metric for model development was maximizing the area-under-curve on the precision-recall graph (AUCPR) for the validation set. Final model convergence was concluded based on the confusion matrix generated (minimized off-diagonal values and maximized main diagonal values) when the model was applied against a test validation data set.

## 3 Results and Discussion

### 3.1 Model generation

The model performance met the intended success criteria given the high proportions of True Positives and True Negatives as shown in Table 1 (main diagonal values). Off-diagonal values (False Positives and False Negatives) are a significant minority bringing in an overall error rate of 4% (bottom right corner).

**Table 1:** The confusion matrix summarizing the validation outcomes for the GBM model.

| PREDICTED | ACTUAL |  |  |  |
| --- | --- | --- | --- | --- |
|  |  | Active | Inactive | Error |
|  | Active | 98% | 2% | 2% |
|  | Inactive | 6% | 94% | 6% |
|  | Totals | 52% | 48% | 4% |

Molecular structure was used as an input to generate both mechanistic and structural factors usable in building the prediction model. They are divisible into these main areas: mass, Hydrophobicity, electro-topological and structural representations. A summary of the descriptor choice is provided in Table 2.

**Table 2:**
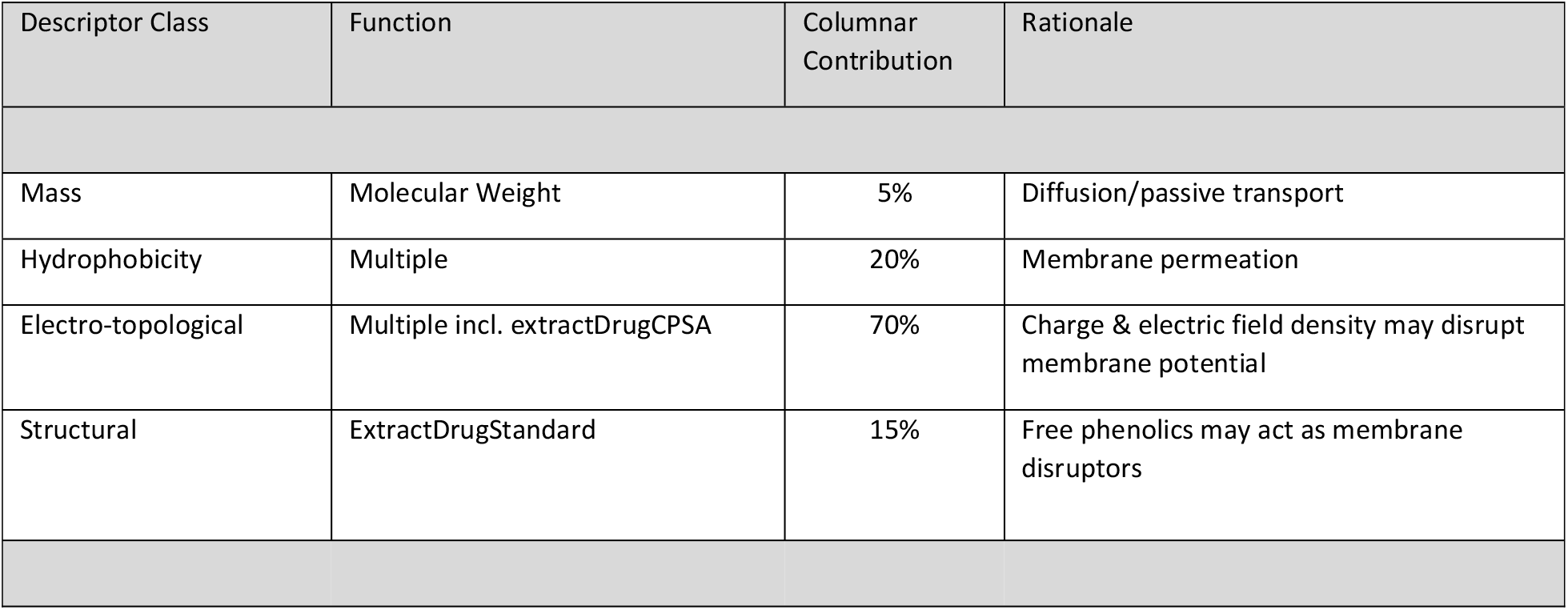
A representative summary of high vaeriance features that contribute significant information to the model.

In Figure 1, the variable importance plot shows the most influential variables in the GBM. There are topological factors (Autocorrelation of a Topological Structure (ATS: ATSp2, ATSp3 & ATSp5, TopoPSA), polarizabilities (bpol, apol & AMR), hydrophobicity (ALogp2), molar mass (MW). These are mechanistic in nature driving the antiplasmodic activity at greater magnitudes than structural features represented in the molecular fingerprints.

**Figure 1:**
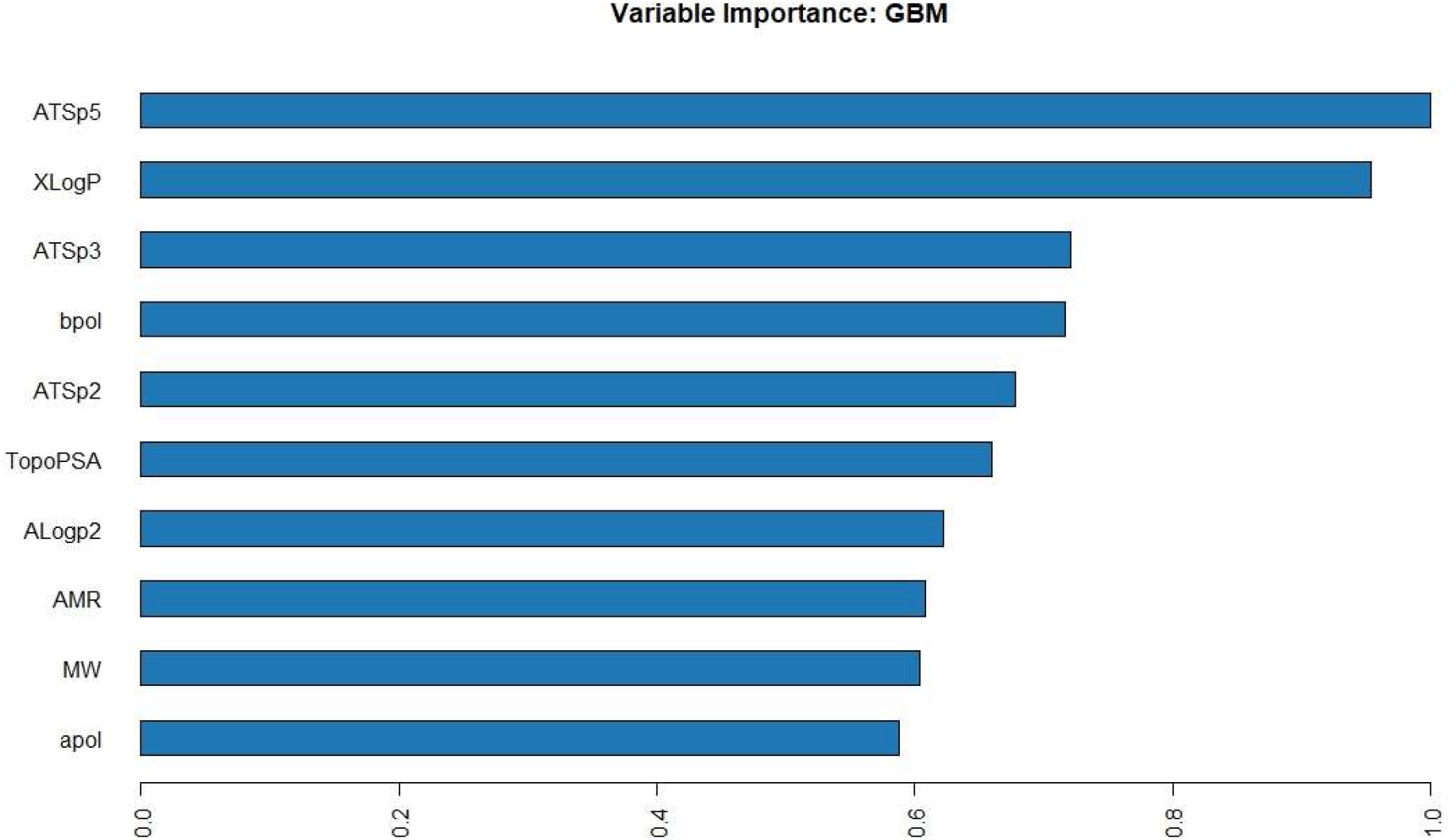
A variable importance plot showing the most influential variables in the GBM; *Validation metrics: AUCPR/AUC (>=0*.*99); Mean Error/Class (0*.*04)*

The contributions in a variable importance plot are given only in absolute terms and for the entire model. However, they do not indicate the individual and directional contributions for each factor one data instance at a time. In Figure 2, SHAP values extracted from members of the validation set show that only structural values have a clear boundary (short simple gradients with negative/red contributions delineated from positive/blue contributions) while electro-topological and hydrophobicity vectors have more complex impact (long varying gradients on the response-vector graphs) on the antiplasmodic activity. This matches the earlier observations where the tail length in the SHAP values correspond to the contributions.

**Figure 2:**
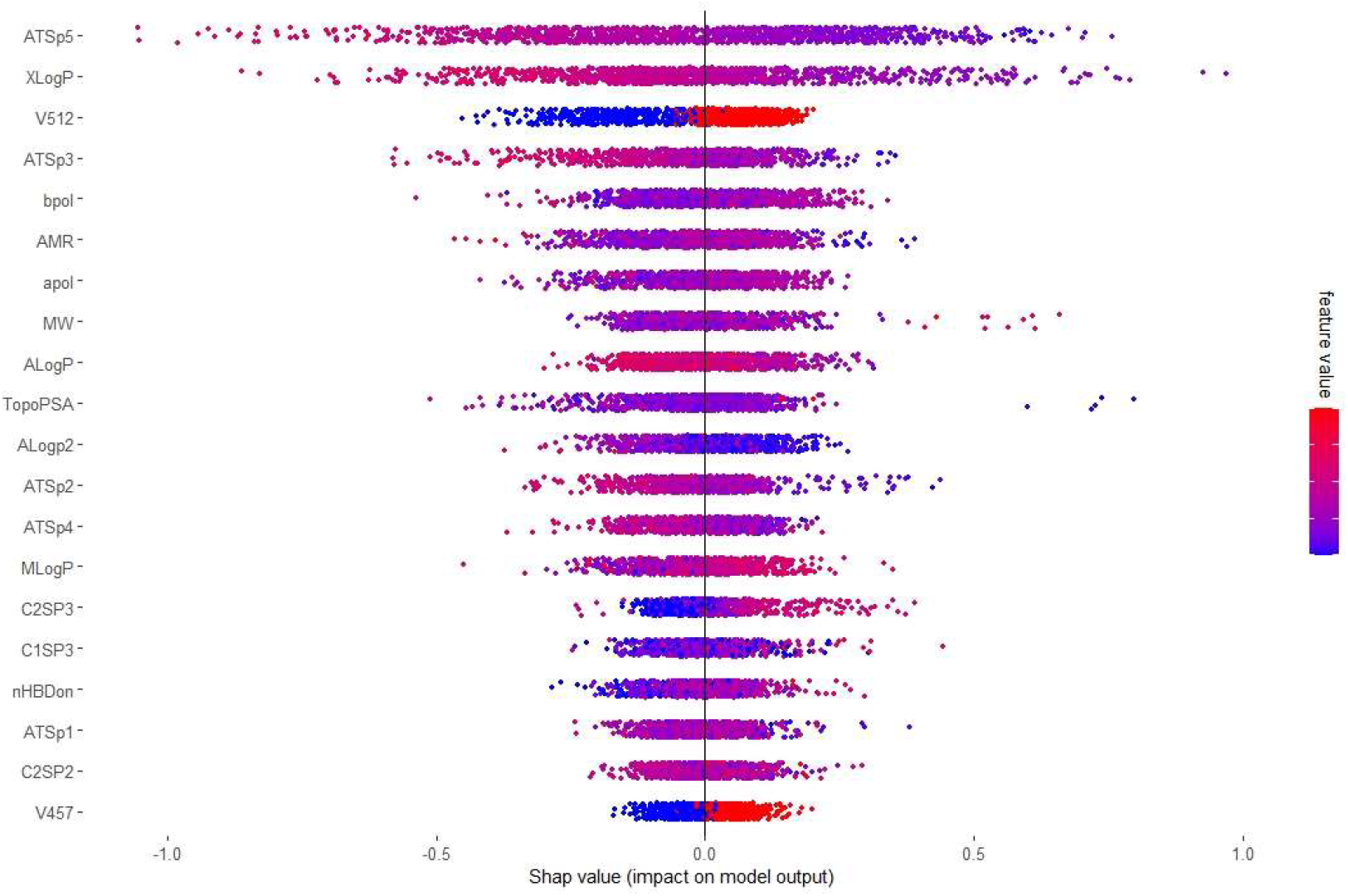
SHAP values applied against the validation set show that only structural values have a clear boundary (simple gradients) while electro-topological and hydrophobicity vectors have more complex impact (varying gradients on the response-vector graphs) on the antiplasmodic activity. *Validation metrics: AUCPR/AUC (>=0*.*99); Mean Error/Class (0*.*04)*

### 3.2 Model Exploration

#### 3.2.1 Hydrophobicity

Hydrophobicity is well known to drive the mechanism by which drugs are shuttled and/or sequestered across membranes and in vesicles. From Figure 3, the x-intercept occurs at around 2.7 which indicates that hydrophobicity is a threshold parameter. Once the threshold is crossed, the parameter contributes positively to the antiplasmodic activity. Notably, the hydrophobicity is calculated from the atomic contribution to the water-octanol partition coefficient.

**Figure 3:**
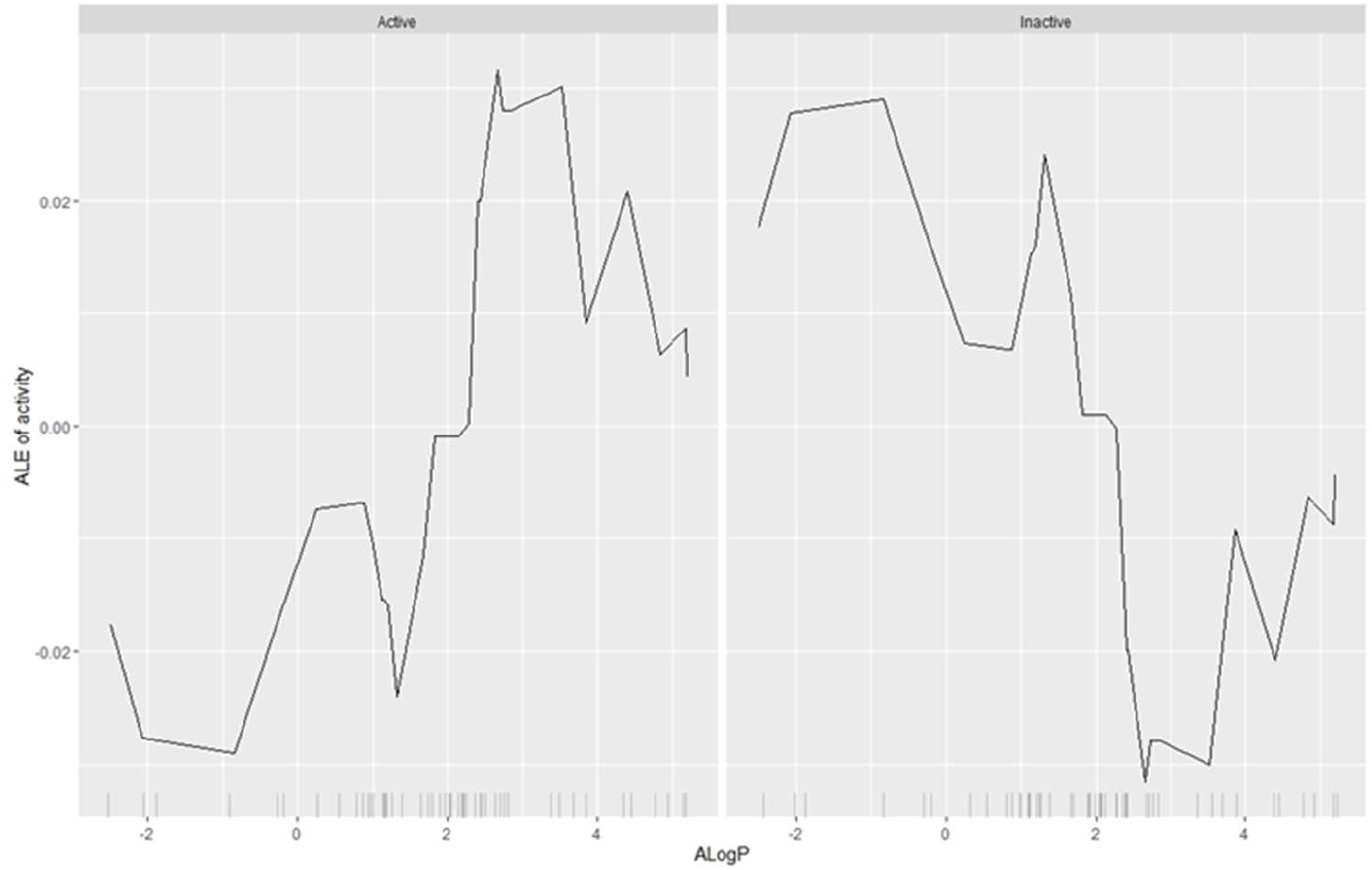
An Accumulated Local Effects (ALE) of the activity plotted against the atomic partition coefficient (AlogP) showing a complex relationship depending on the molecular class and the specific hydrophobicity measure. Generally, increasing hydrophibicity drives antiplasmodic activity.

Additionally, we can examine the hydrophobicity measure using an additivity atomic contribution method trained on a much larger library to improve correction factors. This exercise may help to reinforce previously observed relationships. The threshold of 2.7 seen earlier in Figure 3 also reappears in Figure 4 but with multiple intercepts the earliest of which occurs at 1.8 However, the first meaningful intercept is still at 2.7 since the activity stays at or above the zero line from here onwards.

**Figure 4:**
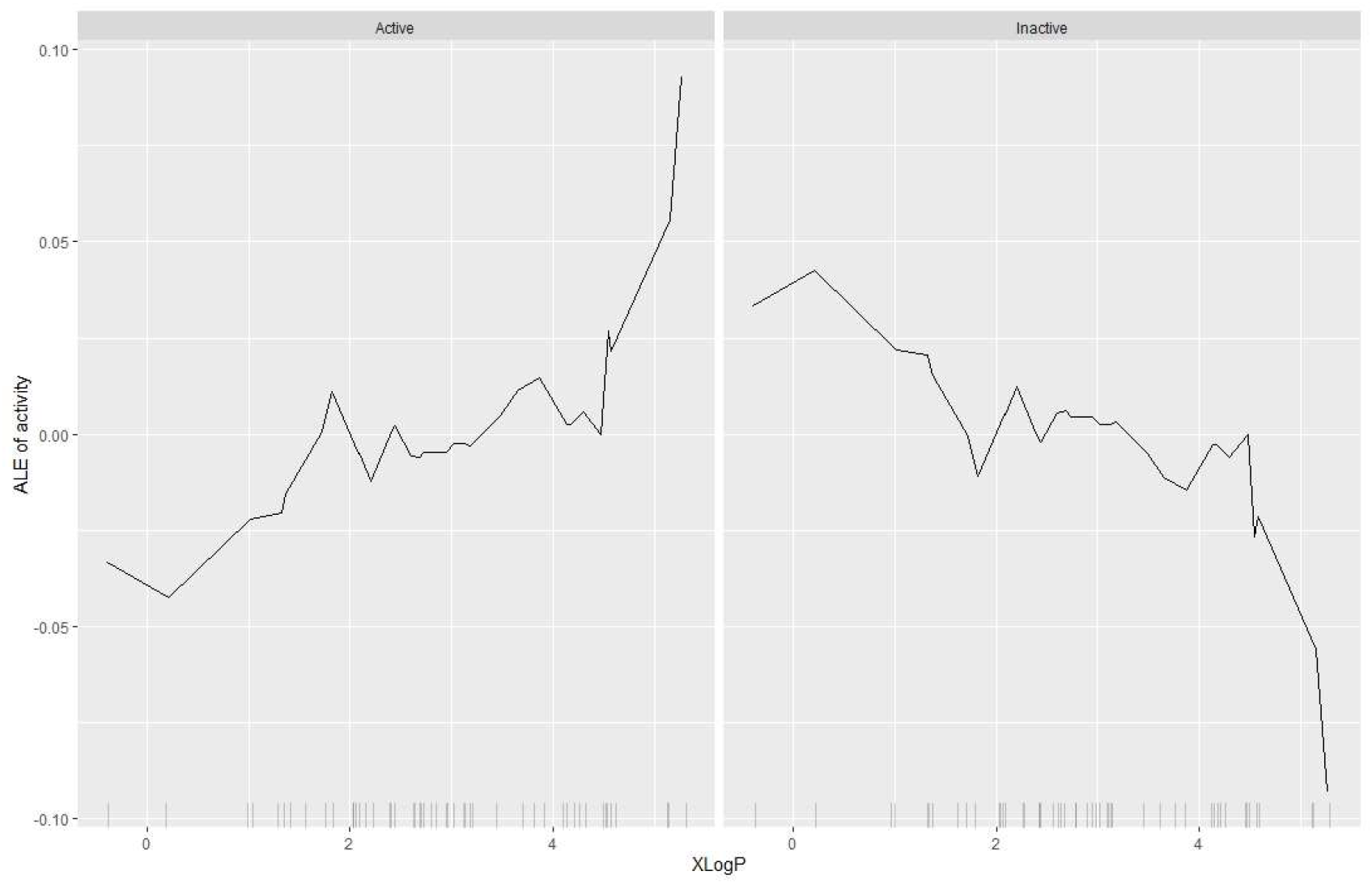
An Accumulated Local Effects (ALE) of the activity plotted against the additive atomic partition coefficient (XLogP) showing a complex relationship depending on the molecular class and the specific hydrophobicity measure. Generally, increasing hydrophobicity drives antiplasmodic activity – this reinforces the relationship observed in Figure 3

#### 3.2.2 Topological Polar Surface Area (TPSA)

The TPSA is well known to affect the ability for molecules to penetrate cell membranes. The TPSA is a sum of all polar surface areas of the molecule with major contributions coming from hydrogen bond donors and acceptors. According to Figure 5, there appears to be a thresholding associated with the effect of TPSA on antiplasmodic activity since the ALE crosses the abscissa reference line (y=0) close to 63 squared angstroms. From this point (x=63 Å^2^) the TPSA has a positive contribution to antiplasmodic activity.

**Figure 5:**
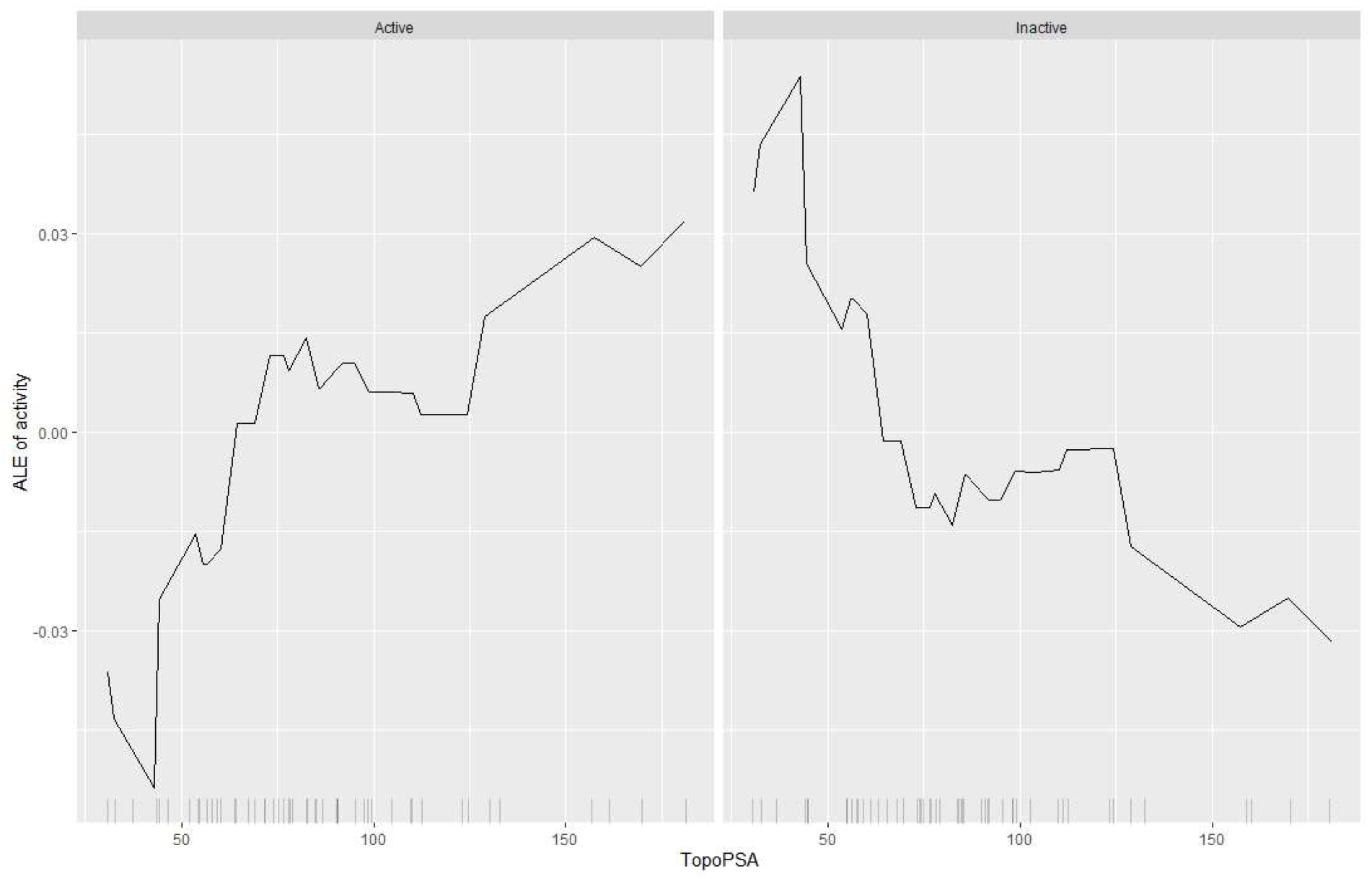
An Accumulated Local Effects (ALE) of the activity plotted against the Topological Polar Surface Area (TPSA) showing a generally increasing activity with increase in the TPSA whose major contributions come from hydrogen bond donors and acceptors. This is mirrored in the inactive class.

#### 3.2.3. Molecular Weight (MW)

Molecular weight affects the passive transport of molecules across membranes. Some rules on drug-likeness reference molecular weight cut offs (MWCO); in this investigation, it appears there is an upper limit just under 500(19). However, it is also clear that antiplasmodic activity being non-linear and multivariate, is a complex relationship of underlying physicochemical parameters. This complexity is captured by the varying activity-MW gradient below the MWCO as shown in Figure 6.

**Figure 6:**
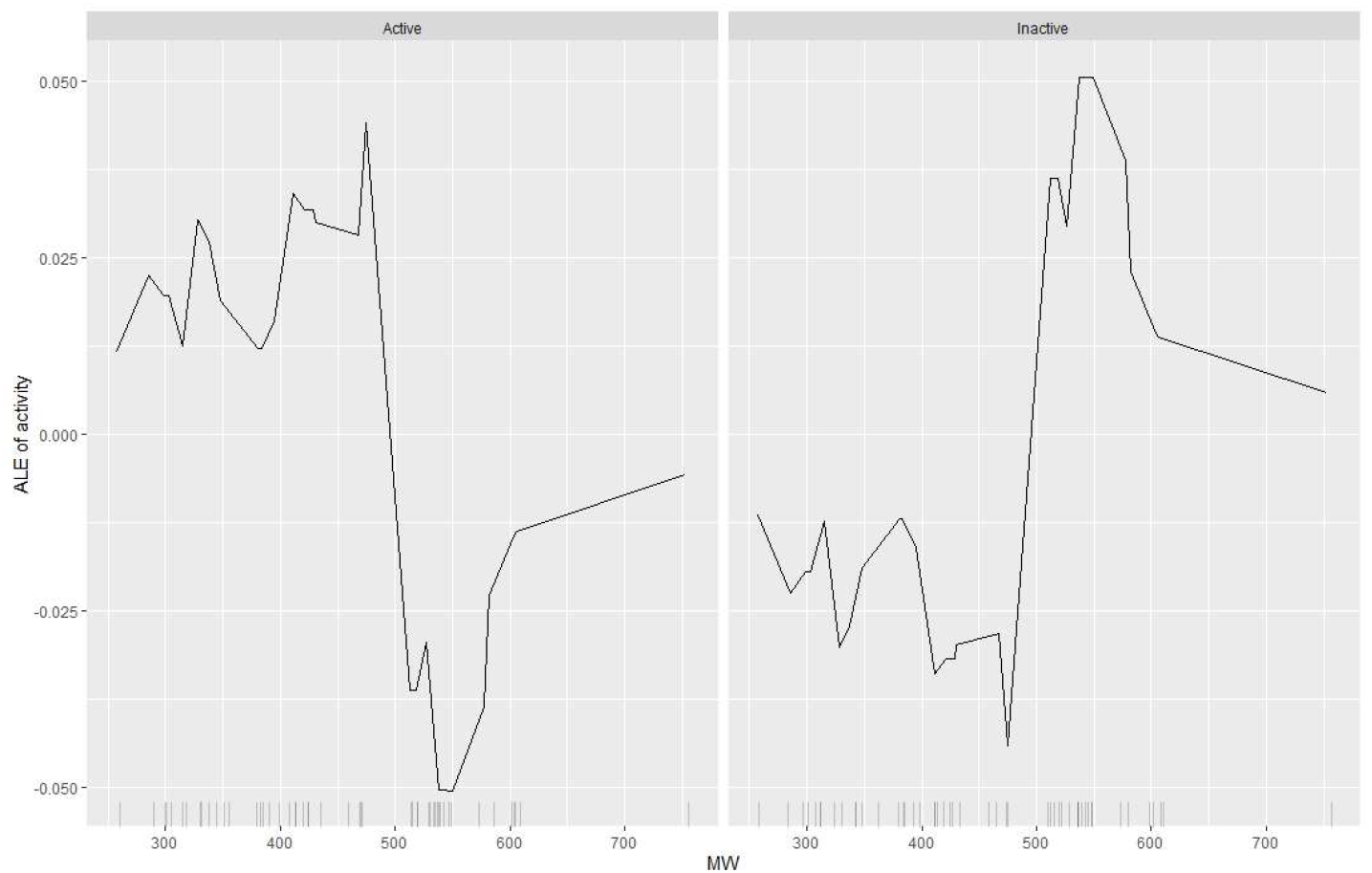
An Accumulated Local Effects (ALE) of the activity plotted against the molecular weight (MW) showing the existence of an MW cut-off. The ALE is mirrored in the *Inactive* class.

#### 3.2.4. Exploring the structural information hidden within the library

The data set is incredibly large and while the classification GBMs are especially helpful in mapping structural and physico-chemical data onto a probabilistic response and defining margins of separations between active and inactive chemistries, the structural meanings behind active molecules are harder to define. A structure-based tree map was therefore extracted from the group of active molecules resulting in a molecular cloud visualization. These representations demonstrated the significance of the thiazole motif (centre, larger) and its structural progenies (highlighted in gold) as shown in Figure 7.

**Figure 7:**
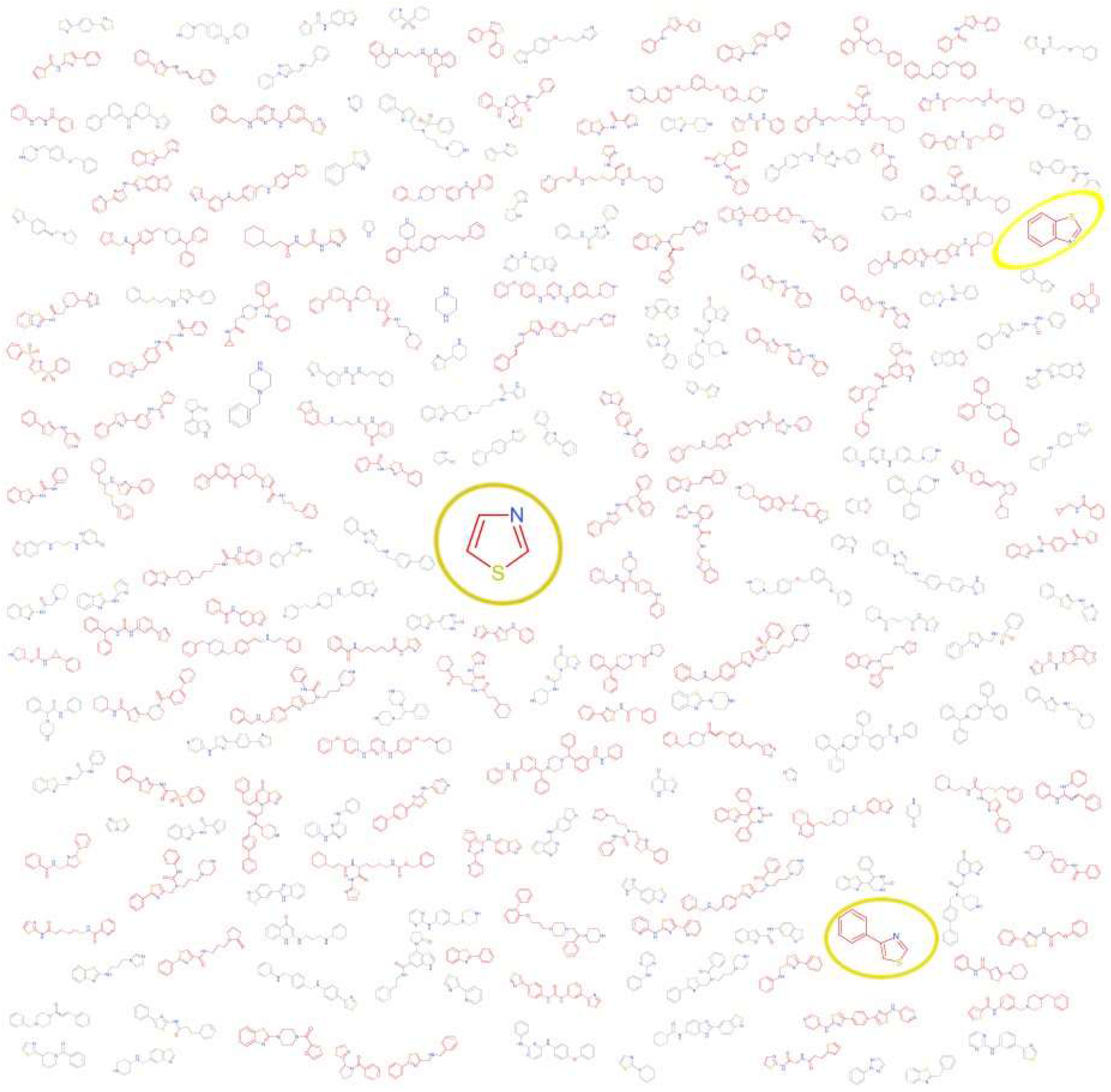
A structure-based tree map extracted from the centre of the molecular cloud visualization. These representations emphasize the significance of the thiadiazole motif (top right, larger) over the diazole motif (bottom right, smaller).

**Figure 8:**
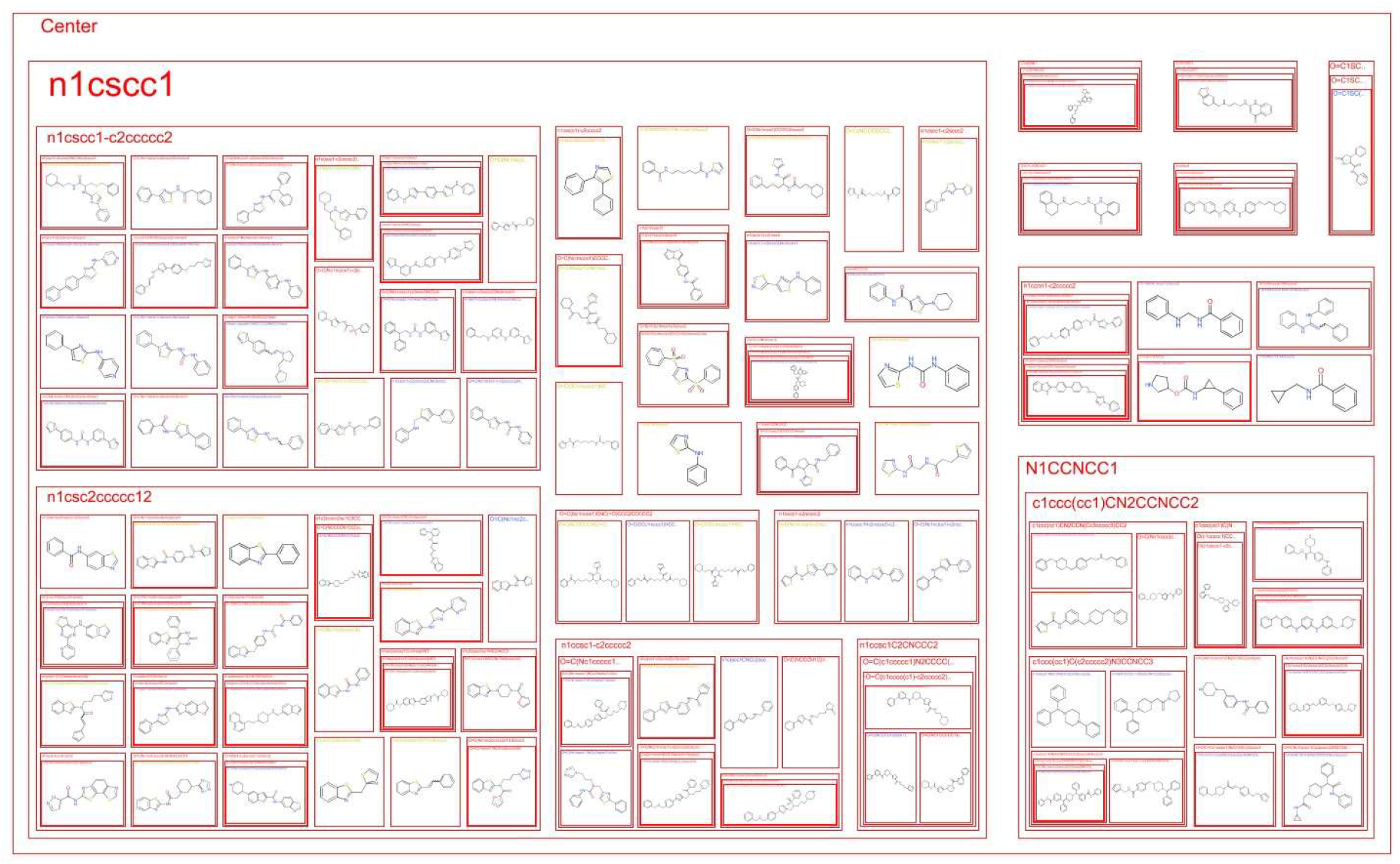
A structure-based tree map extracted from the centre of the molecular cloud visualization. These representations emphasize the significance of the thiazole motif (top right, larger) over the diazole motif (bottom right, smaller).

The structure-based tree map can be extracted from the centre of the molecular cloud visualization. These representations emphasize the significance of the thiazole motif (top right, larger) over the diazole motif (bottom right, smaller). These structural separations are harder to delineate within the full dataset without a visualization tool such as a tree map.

One may also examine the top left (root, Figure 9A) vs. the bottom right (leaf terminus, Figure 9B) to distinguish the left-right ‘poles’ of the tree map. The comparison shows that the root of the molecular map has fewer modified cycloalkanes while the termini have more diazoles (Figure 9B). This demonstrates an increasing aromatic complexity and diversification of the core scaffold.

**Figure 9:**
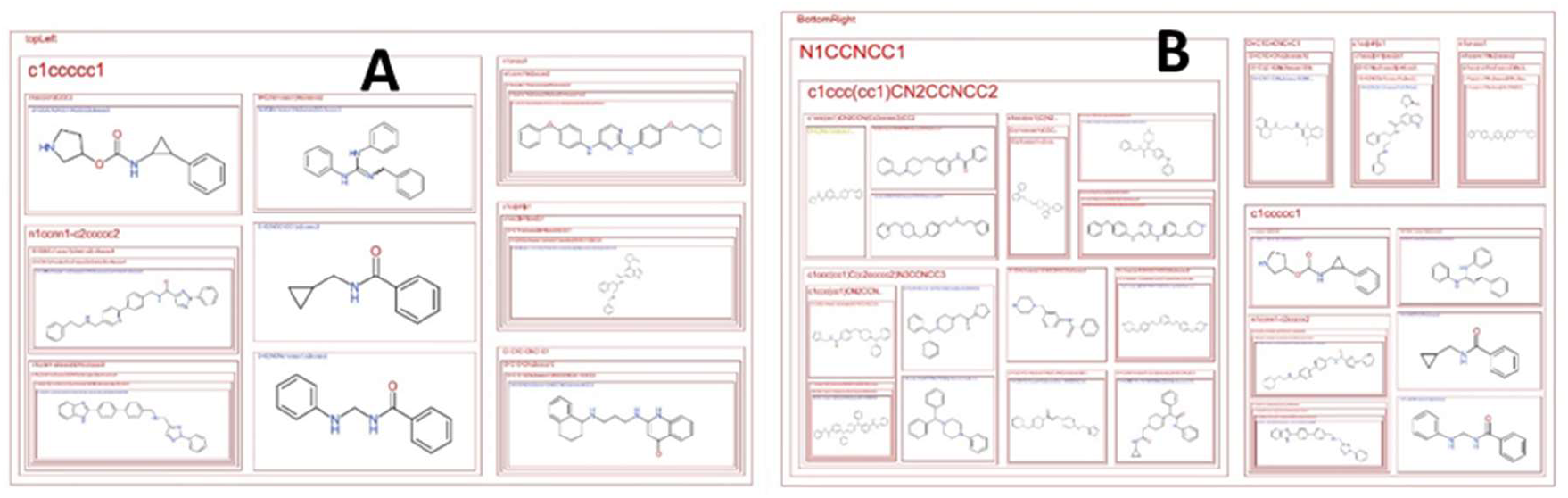
A comparison of the root of the tree map (A) from the leaf (B) shows that the root of the molecular map has fewer modified cycloalkanes while the termini have more diazoles (B).

Examining the central node in the tree map (Figure 10) shows that the two-membered diazole chain is the core scaffold. On the full spectrum of the activity, there are branching extensions to the scaffold resulting in attenuation on the left and sustained activity on the right as shown by the blue-green coloration in a left-right direction.

**Figure 10:**
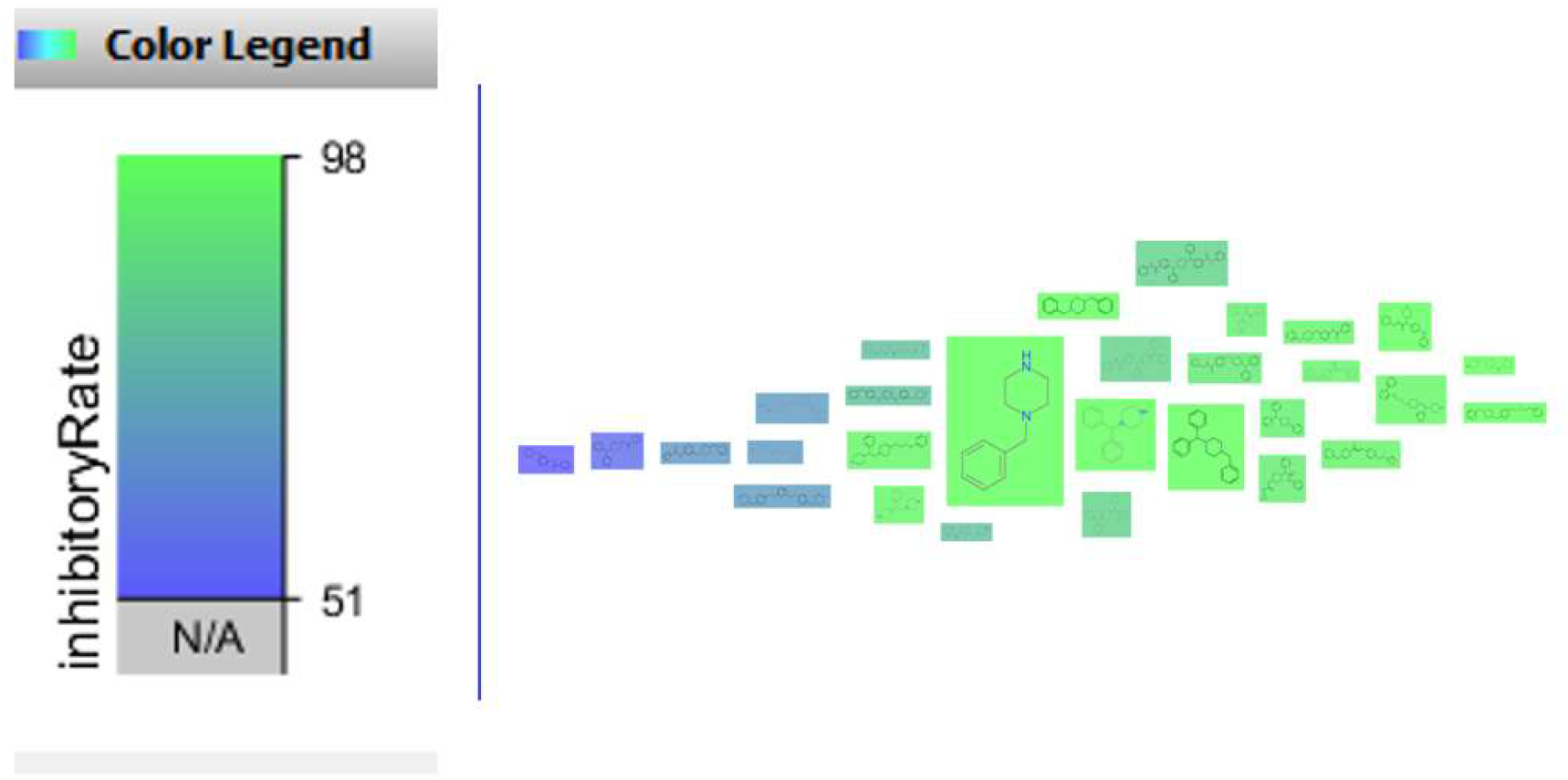
Examining the central portion of the tree map shows there is a full range of inhibitory rates (blue --low while green codes for high inhibition; all molecules display inhibitory activity). Diazoles appear to have a high representation in the molecular cloud.

#### 3.2.5. Case Question: Can molecular redesign from shared core scaffolds identify candidates?

Table 3 shows three molecules of interest with potential antiplasmodial activity. TCMDC is a lead compound that scored strongly in the model training procedures yielding an 89% probability of being active. It is a molecular duplet functioning as a reference molecule. When compared to napthylisoquinolines (Dioncophyline C and Korupensamine E), the standard is a clear positive control. The isoquinoline functionality is present in the positive control and is amplified in the two virtual experiments. However, there are fluoro-groups and ethoxy-or carbonyl linker groups. Generally, the two comparisons are simpler molecules with some core scaffolding shared with the positive control. Mechanistically, the virtual experiments have comparable (in-scope) polarizability, hydrophobicity and charge densities to the control and the full dataset. The virtual experiments have a similarity score of (0.90 – 0.96, MCS range) while the control has a low similarity to the two experiments (0.31 – 0.39) demonstrating significant redesign whilst retaining meaningful antiplasmodic activity.

#### 3.2.6. Case Question: Does bitterness connote antiplasmodic activity?

It has been previously theorized that bitterness is an associated quality to therapeutic value and microbial quorum sensing. Additionally, extreme bitterness may be a connotation for potential toxicity given its observed implication in emesis and learned avoidance. We examined this association by sub-selecting bitter compounds from the library. These are molecules that are at least as bitter as quinine. That is molecules with a Relative Quinine Index of one or less. When the mean RQI for active molecules was compared to that of inactive molecules, the bitterness index was 2.70 times higher for molecules classified as active than for molecules classified as inactive as shown in Table 4.

**Table 3:**
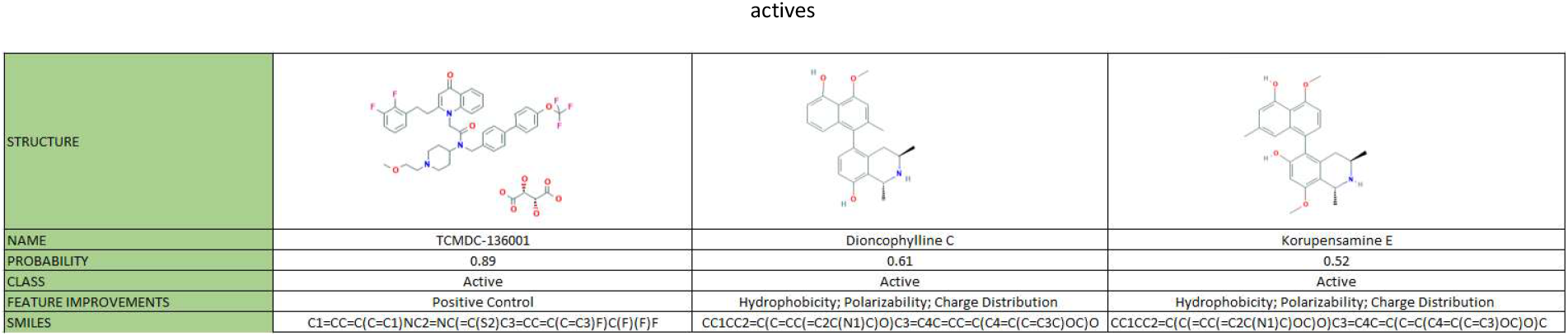
The simulation shows the class of napthylisoquinolines namely Dioncophyline C and Korupensamine E are classified as having antiplasmodial activity but with varying probabilities. In both cases, the hydrophobicity and charge distribution (TPSA) are contributory factors to the positive identification as antiplasmodials actives

**Table 4:**
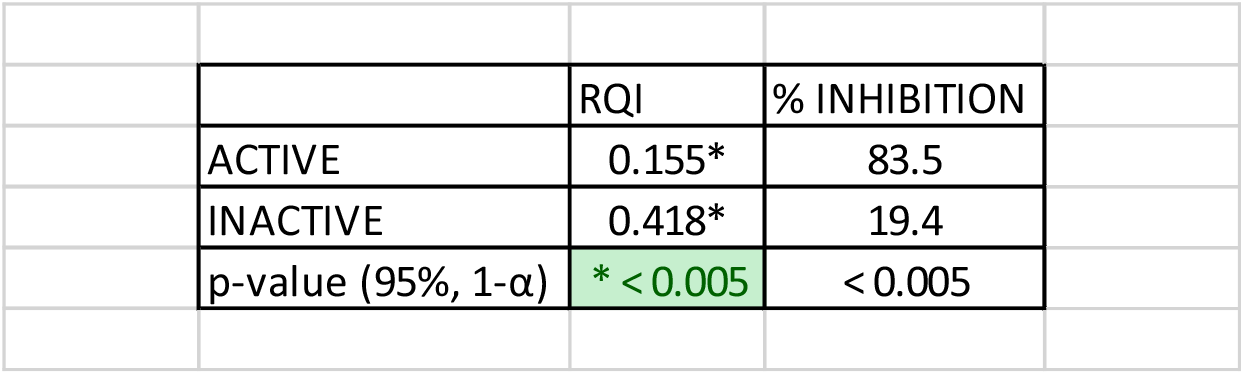
A table showing that for molecules that are at least as bitter as quinine (i.e., having an RQI/Relative Quinine Index of 1 or less), molecules with a high antiplasmodic activity (mean of 83.5%) also have significantly more bitterness than the inactive molecules.

Visualizing the result confirms the significance test with inactive molecules positing a higher RQI than active molecules. This means that inactive molecules are less bitter than the active molecules. While this may pose a challenge for enteric delivery, it can nevertheless be masked when parenteral delivery is not a suitable substitute.

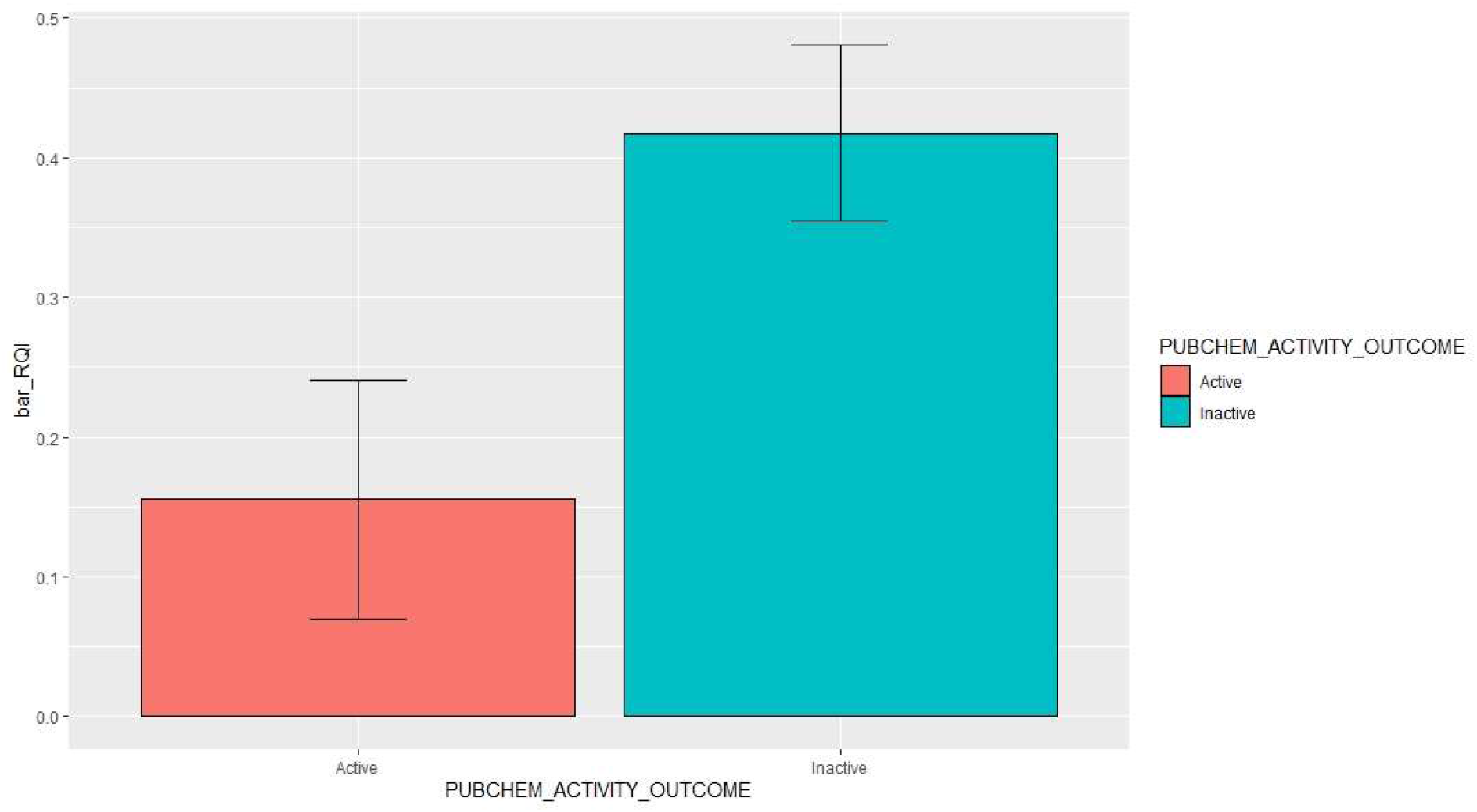

## Conclusion

In conclusion, a GBM-FP-PC approach to classifying small molecules into antiplasmodic classes has been demonstrated to be chemometrically and conceptually valid. By using both mechanistic and fingerprinting input parameters, the model becomes more interpretable in its original form. Furthermore, explanatory models using molecular scaffolding procedures help to uncover human-interpretable explanations for the success of the model. The model may be usable in designing and evaluating small molecules for use in the development of next generation drug design. Further, it may help develop nutritional approaches to supressing parasitaemia in endemic populations when used in conjunction with bitterness models bearing overlapping scope. In this regard, the link between psychophysical senses and antiplasmodic activity can be useful for identifying neighbourhoods in the chemical space that are suitable for molecular design. It is proposed that future work may focus on examining safety and toxicity models to accompany potency predictions. A similar effort behind the identification of durable cocktail formulations that are evolutionarily durable in the fight against malaria will help ensure the effort is beneficial in the long-term. Alternative avenues for research include the examination of chemical adjuvation to reverse plasmodial resistance which may require a corresponding chemical library.

